# Growth arrest is a DNA damage protection strategy in plants

**DOI:** 10.1101/2024.03.12.584648

**Authors:** Antonio Serrano-Mislata, Jorge Hernández-García, Carlos de Ollas, Noel Blanco-Touriñán, Aurelio Gómez-Cadenas, Robert Sablowski, David Alabadí, Miguel A. Blázquez

## Abstract

When exposed to stress, plants slow down their growth while activating defensive mechanisms. This behaviour has been proposed to help plants reallocate resources and meet the energy demands required for survival. In this study, we show instead that plants can grow under limited water availability without compromising their tolerance to the stress. However, cells that continue to divide under stress accumulate DNA damage, which frequently leads to cell death. Given that the DNA lesions are observed in the apical stem cells that give rise to all plant organs, including flowers, we propose that systemic growth arrest is used as a defense strategy that plants employ not only to maximize individual fitness, but also to ensure the accurate transmission of genetic information to their progeny.

## Introduction

The growth-defense trade-off, characterized by the simultaneous restriction of growth and activation of defense mechanisms in response to environmental threats, is observed in diverse organisms, from microbes to plants and mammalian cells ^1–3^. This trade-off is often attributed to competition for shared resources along the growth and defense pathways ^4^. For example, yeast cells prioritize the production of defense proteins by repressing transcripts related to cell division during acute stress ^5^.

Although plants possess cell-autonomous checkpoints for stress-dependent control of cell division that are equivalent to those described in microorganisms and mammalian cell cultures ^6^, the growth-defense trade-off results in a systemic growth arrest. Recent studies challenge the view that the trade-off is caused by simple ‘metabolic competition’ and show that it results from a programmed interaction between cellular signaling pathways ^7,8^. In a first study, the trade-off triggered upon pathogen attack was eliminated by simultaneously enhancing the jasmonate-dependent defense response and reducing the activity of the phytochrome B (phyB)-dependent growth suppression pathway. These plants maintained their robust growth and defense capability against herbivorous insects ^7^. In a second study, the stimulation of a specific branch of brassinosteroid signaling resulted in increased tolerance to drought stress without loss of growth^8^. However, phyB and brassinosteroids regulate multiple aspects of plant physiology. Indeed, loss of phyB activity also resulted in reduced photosynthetic capacity, but the decrease in energy supply was compensated by a reduction in leaf thickness, and brassinosteroids are also known to stimulate plant growth, so a more precise method of uncoupling growth inhibition from defense responses would be required. Moreover, even if the trade-off is not caused by competition for resources, a pivotal question remains unanswered: does the genetically-driven redistribution of resources confer an advantage under stress conditions, or is there an alternative benefit to the observed trade-off? To address this question, we have targeted the DELLA pathway to generate plants that specifically overcome the mechanism of growth inhibition by stress and studied their performance under water limitation. Studies in *Arabidopsis thaliana* have shown that DELLAs play an important role in limiting growth under stress ^9,10^. By eliminating DELLA activity, inflorescence growth can occur even under stress conditions. However, DELLA proteins also promote oxidative stress tolerance ^9^. To separate the role of DELLAs in growth regulation from defense responses, we have focused on the CDK-inhibitors (CKIs) of the KIP-RELATED PROTEIN (KRP) and SIAMESE-RELATED (SMR) families, whose activation by DELLA inhibits cell division ^11,12^. In doing so, we have observed a severe increase in the accumulation of DNA damage linked to cell divisions that could potentially be transmitted to their progeny.

## Results and Discussion

### Loss of *SMR1* suppresses DELLA-mediated growth restriction

Despite the potential redundancy among the members of the KRP and SMR families, *SMR1* emerged as the most relevant DELLA target in the context of inflorescence development, as mutation of *SMR1* alone was sufficient to promote stem elongation in the presence of paclobutrazol (PAC), an inhibitor of gibberellin (GA) biosynthesis that causes DELLA accumulation (Supplementary Fig. 1). The relevance of *SMR1* was confirmed genetically, as only *smr1* suppressed the dwarfism of plants expressing non-degradable versions of the DELLA proteins GAI ^13^ (Fig. 1a; Supplementary Fig. 2a, b) and RGA ^14^ (Supplementary Fig. 2c, d). It is worth noting that *gai-1 smr1* plants developed significantly taller inflorescences and produced three times more reproductive biomass and seeds than *gai-1* (Fig. 1a, b; Supplementary Fig. 3a-c).

**Fig. 1.**
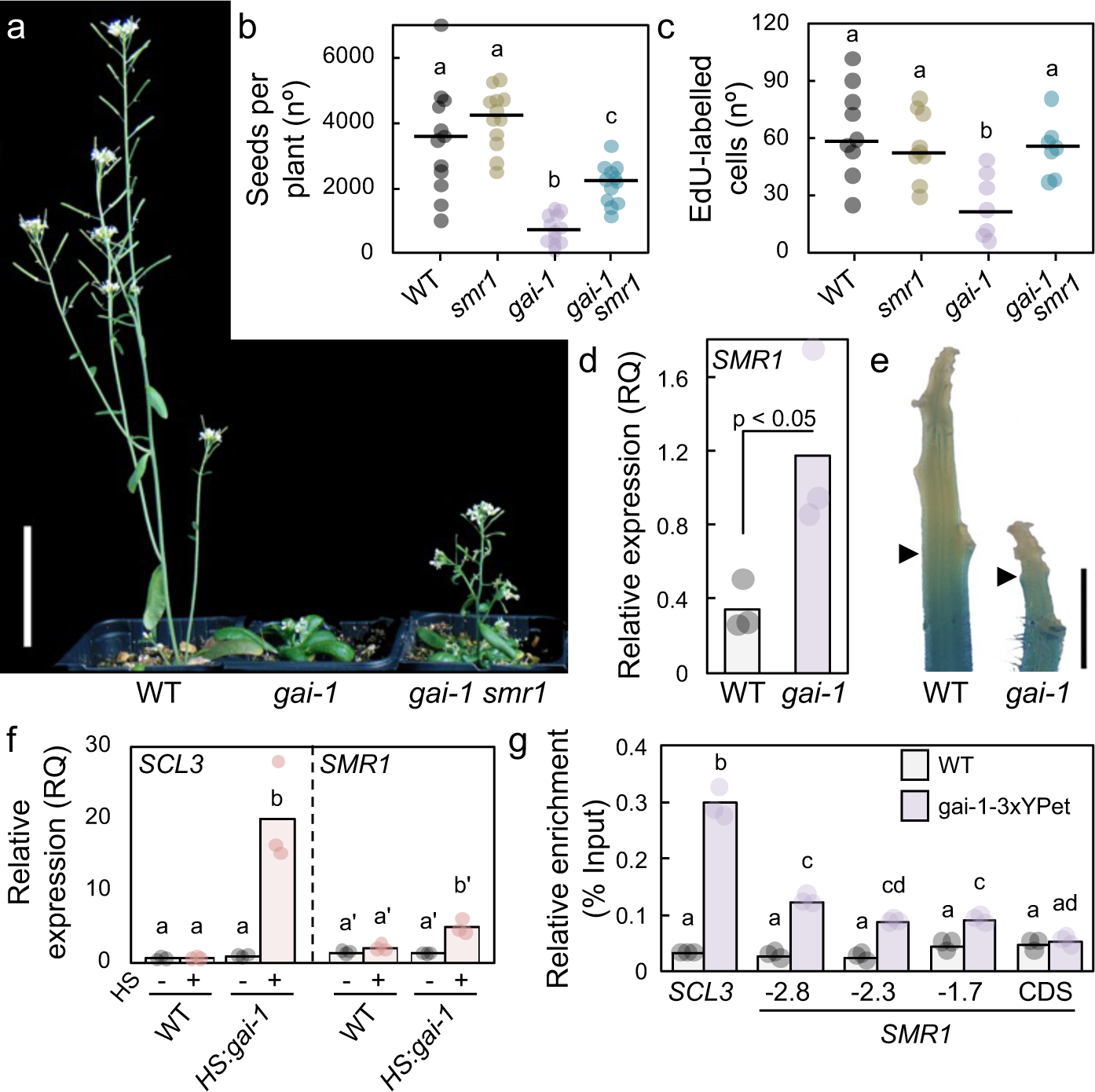
DELLA proteins directly activate *SMR1* transcription to restrict reproductive growth. **a** Representative inflorescences of five weeks-old WT, *gai-1* and *gai-1 smr1* plants. **b-c** Seed production (**b**) and number of EdU-positive cells in the SAM region (**c**) in WT, *smr1*, *gai-1* and *gai-1 smr1* plants. **d** Relative quantity (RQ) of the *SMR1* transcript in dissected 5 mm inflorescence tips. **e** *pSMR1:GUS* reporter activity in representative WT and *gai-1* inflorescence tips. Arrowheads indicate the upper limit of the stained region. **f** Relative quantity of *SCL3* (positive control) and *SMR1* transcripts in WT seedlings and in seedlings expressing a non-degradable GAI protein under the control of a heat-shock inducible promoter *(pHS:gai-1)*. Seedlings were exposed to 37 °C for 30’ (HS+) or left under growth conditions (22 °C, HS-). **g** ChIP-qPCR showing the binding of the gai-1-3xYPet protein to genomic regions upstream the *SMR1* coding sequence (CDS). A fragment of the *SCL3* promoter was amplified as a positive control. WT seedlings were used as negative controls. In (**b**) and (**c**) the dots stand for individual observations [n = 12 in (**b**), n ≥ 7 in (**c**)] and the horizontal lines for mean values. In (**d**), (**f**) and (**g**), bars represent the mean value of three independent biological replicates (represented by dots). In (**b**), (**c**), (**f**) and (**g**) letters indicate statistical groups obtained using Tukey’s post hoc test [p < 0.01 for (**b**), (**c**) and (**g**); p < 0.001 for (**f**)] following one-way ANOVA. In (**d**), *p* value is for the equality of means (two-tailed *t* test). Scale bars, 5 cm in (**a**) and 2 mm in (**e**).

Two observations confirmed that the differences in stem length were due to changes in cell division rates at the shoot apical meristem (SAM). First, *gai-1* mutants had fewer cells labelled with 5-ethynyl-20-deoxyuridine (EdU) in the SAM region than the wild type (WT), which was restored by loss of *SMR1* function (Fig. 1c; Supplementary Fig. 3d). And second, expression of the *pCYCB1;2:GUS* mitotic reporter was reduced in the apical zone of the inflorescence of *gai-1*, as previously reported ^11^, but returned to WT levels by mutating *SMR1* (Supplementary Fig. 3e, f). Importantly, *SMR1* can be considered a direct functional target of DELLA activity for three reasons: its domain of expression was expanded in *gai-1* inflorescence tips (Fig. 1d, e), its expression was rapidly induced following *gai-1* transcriptional activation (Fig. 1f), and the DELLA proteins GAI and RGA were significantly bound to promoter regions of *SMR1* using chromatin immunoprecipitation assays (Fig. 1g; Supplementary Fig. 4). Although stem elongation is still partially impaired in *gai-1 smr1* plants, suggesting that additional factors mediate DELLA inhibition of reproductive growth, our results indicate that mutating *SMR1* is an appropriate strategy to generate plants resistant to DELLA-mediated growth arrest, and that these plants can be used as a tool to test the hypothesis that growth inhibition is critical to guarantee optimal performance under abiotic stress.

### Growth arrest is not required for optimal acclimation to water limitation

DELLA accumulation leads to increased tolerance to different types of abiotic stress, including drought ^15^. Therefore, we investigated the physiological responses of WT, *smr1*, *gai-1* and *gai-1 smr1* plants to decreasing soil moisture. The correlation between leaf relative water content (LRWC, an indicator of plant water status) and soil water potential (SWP, λϑΙ_Soil_, an indicator of soil moisture) was comparable between genotypes (Fig. 2a), demonstrating that all genotypes were being exposed to comparable stress intensities (see Materials and Methods for details). As expected, all plants responded to water limitation with a reduction in transpiration (Fig. 2b). However, transpiration rates normalized per rosette area were higher in *gai-1* and *gai-1 smr1* plants under stress (Fig. 2c), which could be explained by the higher stomatal density in *gai-1* leaves ^16^. Despite the higher water loss, the capacity of *gai-1* plants to cope with water limitation in survival assays was better than that of WT plants (Fig. 2d, e). It is likely that this capacity is related to DELLA-enhanced production of antioxidants that buffer the toxic effect of reactive oxygen species (ROS) generated during extreme drought ^17–19^. In agreement with this idea, we observed an increased expression of genes involved in flavonol biosynthesis in stressed *gai-1* plants (Supplementary Fig. 5a, b), accompanied by a higher accumulation of flavonols (Supplementary Fig. 5c) and a reduction of oxidative damage in root cells (Supplementary Fig. 5d, e).

**Fig. 2.**
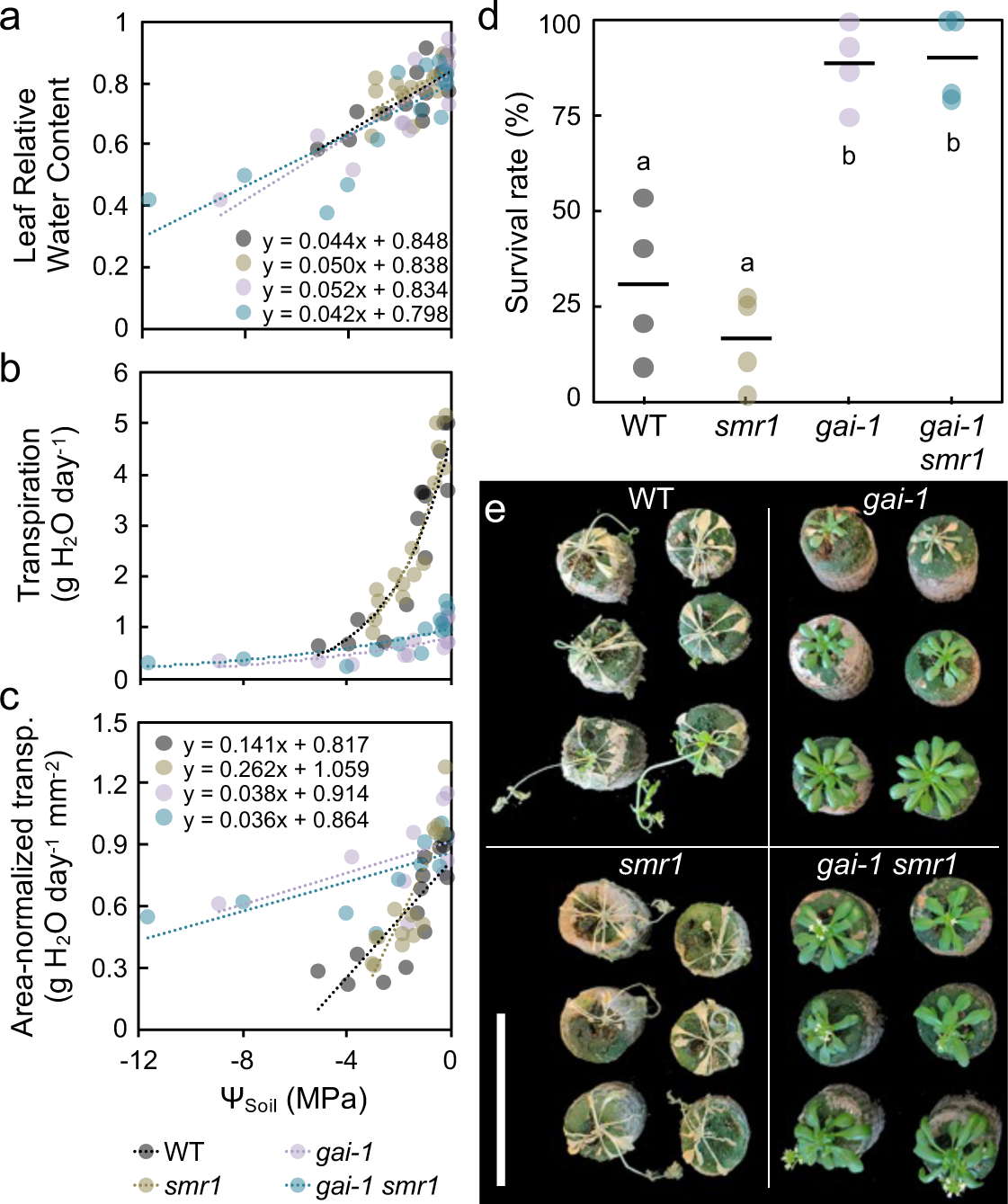
*smr1* has no effect on the drought tolerance of *gai-1* mutants. **a-c** Relative water content in rosette leaves (LRWC) (**a**), daily plant transpiration (**b**) and transpiration normalized to the rosette area (**c**) as a function of soil water potential (ψ_soil_) in WT, *smr1*, *gai-1* and *gai-1 smr1* plants. The dots denote individual plants (n ≥ 15) and the dashed lines represent the fitting functions of the data for each genotype. **d-e** Drought resistance of WT, *smr1*, *gai-1* and *gai-1 smr1.* Four-weeks old plants were maintained at 15-20% LRWC for four days and survival rates were determined three days after rewatering. In (**d**), the dots represent independent assays, each comprising 8-16 plants per genotype. The black horizontal lines represent the mean values. Letters indicate statistical groups obtained using Tukey’s post hoc test (p < 0.001) following one-way ANOVA. In (**e**), representative images after rewatering the plants are shown. In contrast to *gai-1*, the inflorescence of *gai-1 smr1* plants had bolted and contained mature flowers and fruits by the end of the assay. Scale bar, 10 cm.

More importantly, the antioxidant protection and survival rate of *gai-1 smr1* plants was the same as that of *gai-1* plants (Fig. 2d, e), and an equivalent behavior was observed when comparing *rga-τ<17 smr1* and *rga-τ<17* plants (Supplementary Fig. 5f). These data suggest that maintaining increased growth and biomass production does not compromise the proper acclimation and survival of plants exposed to drought stress.

### Cell divisions aggravate genotoxicity

If the energy demands of the defense mechanisms do not justify the need to arrest growth, what is the reason for the conservation of this physiological response in plants over evolution? One possibility is that growth arrest is a tolerated consequence of a more valuable response: the DELLA-dependent stimulation of the response against oxidative stress ^9^. However, this is unlikely because there are additional DELLA-independent mechanisms that promote stress-induced growth arrest ^20,21^. A second possibility is that growth cessation due to cell cycle arrest is intrinsically beneficial under stress. In fact, DNA replication during the S phase of the cell cycle represents a vulnerable point for DNA damage, due to the open chromatin conformation ^22,23^. In concert with this, roots incubated with zeocin, a radiomimetic drug that induces double-strand breaks in DNA, caused increased cell death that was more evident in the dividing cells of the meristem than in the elongation and differentiation zones (Fig. 3a; Supplementary Fig 6). This effect was further enhanced when cell divisions were stimulated by the presence of sucrose in the medium (Supplementary Fig. 6a-c) or by *della* loss-of-function (Supplementary Fig. 6d-f).

**Fig. 3.**
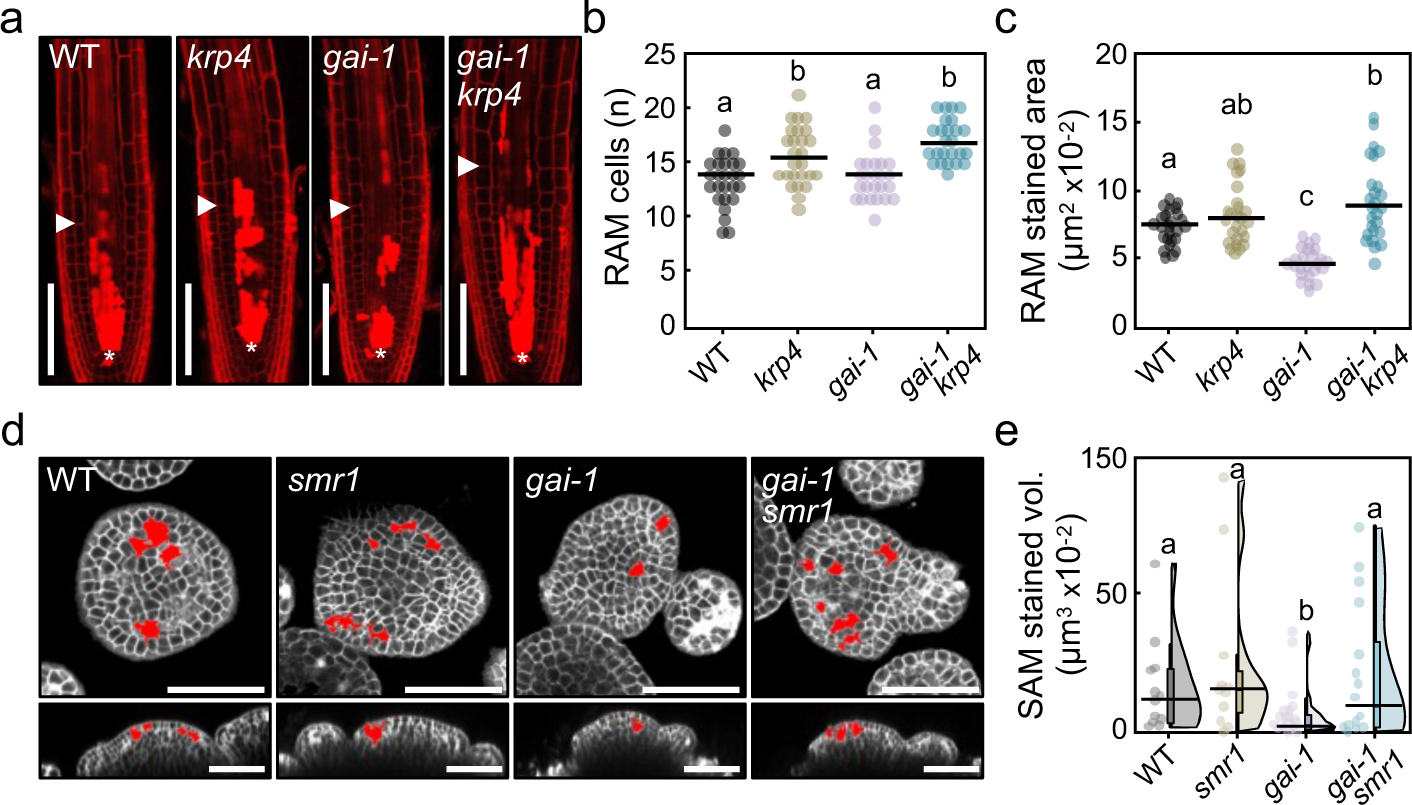
Cell divisions overcome DELLA protection against genotoxic stress. **a** Representative confocal images of WT, *krp4*, *gai-1* and *gai krp4* root tips treated with 20 μg/ml zeocin for 24 h and stained with PI, which colours cell borders but penetrates dead cells. The asterisks mark the position of the quiescent center and the arrowheads mark the RAM length. **b-c** Number of cells in the cortex layer (**b**) and quantification of cell death (**c**) in longitudinal sections through the center of the RAM of zeocin-treated seedlings. The dots represent individual values and the horizontal lines represent the mean value per genotype, n > 25. **d** Orthogonal views of confocal image stacks of representative WT, *smr1*, *gai-1* and *gai-1 smr1* inflorescence apices treated with 100 μg/ml zeocin for 48 h and stained with PI. Dead cells within the SAM region are false colored in red. **e** Cell death volumes in the SAM region of zeocin-treated inflorescence tips. Raincloud plots show the raw data (n ≥ 9, dots), the median value for each genotype (black horizontal lines) and the distribution of the data (boxplots and split-half violin plots). Letters denote statistical groups defined by Tukey’s post hoc test following one-way ANOVA [p < 0.05 for (**b**) and p < 0.01 for (**c**)] or by Dunn’s post hoc test following multiple comparisons Kruskal-Wallis [p < 0.05 for (**e**)]. Scale bars, 50 μm.

The necessity of DELLA for protection against genotoxic agents could be explained either by the prevention of DNA damage by reducing cell divisions or by DELLA-dependent promotion of the cellular DNA Damage Response (DDR) pathway ^24^, for which there is no evidence to date. That the observed genotoxicity is related to cell division capacity was confirmed by analyzing zeocin-induced cell death in the roots of plants with mutant CKIs. Contrary to shoots, SMR1 is hardly relevant for cell divisions in roots (Supplementary Fig. 6i-l), where KRP4 is a more important CKI ^25^. Root meristems of *krp4* and *gai-1 krp4* mutants were larger than those of the corresponding controls (Fig. 3a, b), and the sensitivity to zeocin was higher in *gai-1 krp4* than in the *gai-1* mutant (Fig. 3a, c). Similarly, limiting cell divisions by the activity of DELLA in the SAM resulted in increased resistance to zeocin (Fig. 3d, e; Supplementary Fig. 7a-c), and this effect was reversed by *smr1* (Fig. 3d, e), suggesting that protection of DELLA against genotoxic stress can be abolished by stimulating cell proliferation. Two observations provide additional evidence for the correlation between cell divisions and genotoxicity in the shoot apex. First, the analysis of a *pCLV3:GFP(-ER)* stem-cell niche reporter line showed that the early progeny of stem cells were more sensitive to the drug (Supplementary Fig. 7d). Second, the excessive proliferation of SAM cells in *clavata1* (*clv1*) mutants with impaired stem cell homeostasis ^26^ also resulted in larger regions of cell-death following zeocin exposure (Supplementary Fig. 7e-f). We conclude that limiting cell divisions protects against genotoxic stress in plant meristems.

### Growth arrest under drought protects genome integrity

Environmental stress factors, such as drought, have been shown to increase the generation of ROS ^17^, thereby compromising genome integrity ^27^. To investigate whether plants under stress stop growing to avoid DNA damage, we examined the activity of SAMs under drought and determined the extent of DNA damage and cell death. Drought restricts overall plant growth and the SAM is arrested when the LRWC falls below 50% ^28,29^. We cultivated WT and *della^KO^* plants with sufficient water until flowering and then gradually reduced soil moisture to lower the LRWC to 40-50%, which occurred 12 days later in both genotypes (Fig. 4a; Supplementary Fig. 8a). While water limitation restricted inflorescence growth and stopped the production of new flowers in the WT plants, the SAM of the *della^KO^* mutants remained active and produced new flowers even at LRWC below 50%, although at a lower rate than under well-watered conditions (Fig 4a; Supplementary Fig. 8a-c).

**Fig. 4.**
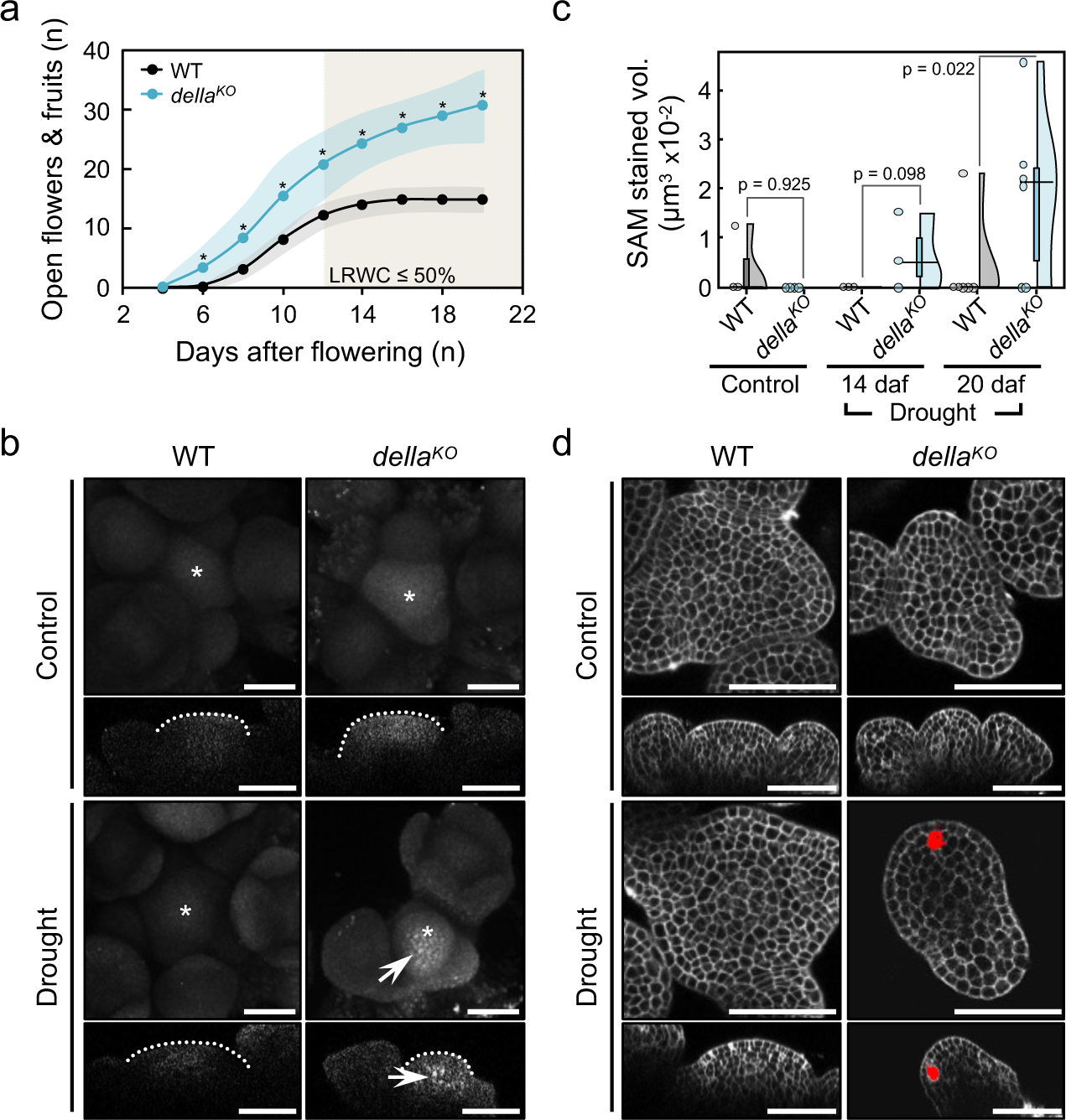
SAM arrest protects DNA from drought stress. **a** Cumulative number of mature flowers and fruits in the main inflorescence of WT and *della^KO^*plants exposed to controlled water restriction after flowering (see Supplementary Fig. 8b for flowers and fruits production in control plants grown under well-watered conditions). The graph shows the mean (dots) and standard deviation (shading) of 12-15 plants per genotype. Asterisks indicate statistical differences, p < 0.01 (two-tailed *t* test). LRWC, relative water content in leaves (see Supplementary Fig. 8a for values). **b** TUNEL labelling to detect DNA breaks in the shoot apex of WT and *della^KO^* plants grown under well-watered (Control) or restricted watering (Drought) conditions for 20 days after flowering. For each genotype and condition, confocal maximum intensity projections are shown in the top panels (asterisks mark the center of the SAM) and the corresponding longitudinal sections are shown in the bottom panels (dashed lines draw the SAM boundary). TUNEL fluorescence was only detected in cells of the SAM of *della^KO^* mutants subjected to drought stress (white signal, marked with arrows). **c** Quantification of cell death in the SAM region of WT and *della^KO^* plants grown under different watering regimes. Control, well-watered plants at a similar stage of development, determined as the number of mature flowers in the main inflorescence, than drought-stressed plants. Drought, plants with limited watering for the indicated number of days after flowering (daf). Raincloud plots show independent observations (n = 3-4 for control and 14 daf plants, n = 5-6 for 20 daf plants), median values (black horizontal lines) and distribution of data (boxplots and split-half violin plots). *p* values correspond to one-tailed Mann-Whitney U tests. **d** PI staining to detect cell death (false coloured in red) in the SAM region of WT and *della^KO^* plants grown under control conditions or drought stress. Orthogonal views of confocal image stacks of representative shoot apices are shown. Scale bars, 50 μm.

Importantly, we found that the more active *della^KO^* SAMs also accumulated higher levels of DNA damage under drought than WT, determined by terminal deoxynucleotidyl transferase-mediated dUTP nick-end labelling (TUNEL) ^30^ (Fig. 4b; Supplementary Fig. 8d). Consistent with the increased accumulation of DNA damage, cell death patches were larger and more frequent in the *della^KO^* SAM than in the WT SAM subjected to drought stress (Fig. 4c, d; Supplementary Fig. 8e).

A reduction in growth rate has been shown to be critical for survival under stress in animal cells in culture ^31^ and in yeast ^32^, which is associated with a reallocation of resources towards functions of stress tolerance ^33^. In contrast, we have shown that efficient defense can occur in plants even when this redistribution is limited, perhaps because plant growth is mainly governed by a small group of dividing stem cells at the meristems that give rise to vegetative and reproductive organs. Growth arrest specifically at the meristems appears to be a critical strategy for plant survival, as these cells are hypersensitive to environmental threats. In barley, drought conditions that minimally affect other organs inhibit the formation of new primordia by the SAM ^34^. Likewise, exposure to low temperatures induces cell-cycle arrest in the SAM region and in the youngest leaf primordia, with minimal impact on DNA replication in older leaves ^35^.

Additionally, in line with the role of the SAM in progeny generation, the inner layers constituting the germline are shielded from DNA damage by localized expression of genes involved in DDR and telomere maintenance in L2 and L3, but not in L1 ^36,37^. The extensive DNA damage (Fig. 4b) and cell death (Fig 3 and 4c, d) observed under stress is therefore expected to result in germline mutations. Stress has been shown to accelerate mutation rates in plants, which are efficiently transmitted to the progeny ^38,39^. This suggests that while growth arrest under stress may be an evolutionarily ancient cell-autonomous response, it has been engineered in plants to trigger a systemic arrest of an entire individual to safeguard not only its own genomic integrity, but potentially that of its offspring.

## Methods

### Plant material

The *Arabidopsis thaliana* Landsberg *erecta* (L*er*) and Columbia (Col) accessions were used as wild-type controls. The following lines in the L*er* background have been described elsewhere: *della^KO^*, *gai-1*, *pRGA:GFP-rga-Δ17*, *pGAI:gai-1-3xYPet*, *smr1-1*, *krp2-3*, *krp4-2*, *clv1-4* and *pCLV3:GFP(-ER)* ^11,14,26,40–45^. The following lines in the Col background were also described: *pHS:gai-1*, *sim-1*, *smr2-1* and *krp1-1* ^46–49^. The double mutant lines *gai-1 smr1-1*, *gai-1 krp2-3*, *gai-1 krp4-2* and *pRGA:GFP-rga-Δ17 smr1-1* were generated by crossing. The *pCYCB1;2:GUS* reporter line, originally in the Col background ^50^, was backcrossed three times with L*er* before being introduced into *gai-1* and *gai-1 smr1-1* mutants. Mutant combinations were identified by PCR with the primers listed in Supplementary Table 1.

### Growth conditions

Arabidopsis seeds were sown on a mixture of sphagnum:perlite:vermiculite (2:1:1) or on 41 mm Jiffy peat plugs (Semillas Batlle S.A., Barcelona, Spain) for experiments that required tight control of watering. After stratification at 4 °C for 5-7 days, the plants were grown under controlled conditions of temperature (22 °C), photoperiod (16/8 h light/dark), and light intensity (100 μmol m^-2^ s^-1^). Transpiration and drought survival assays were performed at 23/18 °C day/night temperature, 8/16 h light/dark photoperiod, 100 μmol m^-2^ s^-1^ light intensity and 60-65% humidity. For the in vitro experiments, seeds were surface sterilized and sown on horizontal or vertical plates containing half-strength Murashige and Skoog Medium including Vitamins (½ MS, Duchefa) with 0.8% phytoagar and 1% sucrose, pH 5.7. After stratification, the plates were placed in cabinets with a 16 h light photoperiod at 22 °C, unless otherwise stated.

### Transpiration and drought survival assays

To determine transpiration rates, plants were grown on Jiffy peat plugs covered with plastic shells (pots) to minimize substrate evaporation. Four weeks-old plants were randomly divided into four groups of four plants each and differentially watered for ten days to achieve a range of soil water potentials from well-watered conditions to plant wilting (0 to −12 MPa) ^51,52^. The pots were then weighed at 24 h intervals to account for transpiration water loss in the plants. Projected rosette areas were determined to normalize whole-plant transpiration per mm^2^. To monitor the relative leaf water content (LRWC) of individual plants, fully developed rosette leaves were cut, weighed (fresh weight) and placed in 2 ml plastic tubes filled with tap water. After 24 h at 4 °C, the leaves were weighed again (saturation weight) and the LRWC was calculated as a quotient between fresh weight and saturation weight. Finally, rosettes were harvested to quantify flavonols and analyse gene expression, as described below.

For the survival assays, four weeks-old plants growing on covered peat plugs were watered to reach the same low soil water content (20% volumetric water content), which corresponds to a point close to a sharp drop in the soil water potential ^52^. Once the soil water content of all plants was homogenised, watering was stopped. Four days after water withdrawal, the plants were watered again until saturation and checked for survival three days later.

### Generation of a *pSMR1:GUS* reporter line

The *pSMR1:GUS* construct contained the *β-glucuronidase* gene (*GUS*) flanked by 3845 bp and 3715 bp of the 5’ and 3’ genomic regions between *SMR1* and the flanking genes. The 5’ region was amplified from genomic DNA of the L*er* ecotype with the primers SMR1 5’ F and SMR1 5’ R (Supplementary Table 1). This fragment was cloned into the pDONR221 vector using Gateway BP recombination (Thermo Fisher Scientific) to generate pENTR-5’gSMR1. The 3’ region was obtained from L*er* genomic DNA in two independent reactions. A 2.3 kb fragment from the *SMR1* STOP codon to a *Hind*III restriction site 2264 bb downstream (3’ fragment 1, f1) was amplified with the primers SMR1 3’ f1 F and SMR1 3’ f1 R (Supplementary Table 1). A 1.4 kb fragment from the *Hind*III site to the downstream gene (3’ fragment 2, f2) was amplified with the primers SMR1 3’ f2 F and SMR1 3’ f2 R (Supplementary Table 1). After A-tailing, f1 and f2 were cloned into the pGEM-T Easy vector (Promega) to generate pGEMT-3’gSMR1-f1 and pGEMT-3’gSMR1-f2, respectively. Both genomic fragments were fused at the *Hind*III/*Pst*I sites of pGEMT-3’gSMR1-f1 to produce pGEMT-3’gSMR1. To generate the *pSMR1:GUS* reporter construct, the 5’ genomic region of pENTR-5’gSMR1 was cloned upstream of the *GUS* gene into the pMpGWB104 binary vector ^53^ using Gateway LR recombination (Thermo Fisher Scientific). The 3’ genomic region of pGEMT-3’gSMR1 was cloned into the resulting vector as *Asc*I/*Sac*I, replacing the nopaline synthase terminator. The *pSMR1:GUS* plasmid was introduced into WT plants of the L*er* ecotype by floral dip ^54^. A representative homozygous line carrying the transgene at a single locus was selected for analysis.

### GUS staining

Shoot apices at a similar developmental stage (with one to three open flowers) were pre-fixed in 90% cold acetone for 20 min, briefly washed with staining solution without X-Gluc and vacuum-infiltrated in staining solution for 10 min. For *SMR1p:GUS* samples, the staining solution contained 50 mM sodium phosphate buffer (pH 7.2), 2 mM potassium ferrocyanide, 2 mM potassium ferricyanide, 0.2% Triton X-100 and 1 mM 5-bromo-4-chloro-3-indolylb-D-glucuronic acid (X-Gluc). For *CYCB1;2p:GUS* samples, with weaker GUS activity, the staining solution did not contain ferro/ferricyanide and the X-Gluc concentration was increased to 2 mM. After vacuum infiltration, the apices were incubated at 37 °C for 18 *(SMR1p:GUS)* or 48 h *(CYCB1;2p:GUS)*. The samples were de-stained with increasing concentrations of ethanol (30, 50 and 70%) and stored in 70% ethanol at 4 °C. Images of representative apices were obtained using a Leica DMS1000 digital microscope.

### Quantification of flavonols

Quercetin and kaempferol content were measured in the rosettes of WT, *smr1*, *gai-1* and *gai-1 smr1* plants exposed to a range of soil water potentials from −0.15 to −4 MPa (from well-watered to moderate-strong drought conditions). Five to six individuals were analyzed for each genotype. Flavonols were extracted in 75% acetonitrile/water supplemented with 1 ppm genistein. To release the aglicones, the extract was mixed with an equal volume of HCl 2N and incubated for 2 h at 80 °C. 1 μl was injected into an Orbitrap Exploris 120 mass spectrometer coupled to a Vanquish UHPLC system (Thermo Fisher Scientific, Waltham, MA, USA). Reverse-phase ultraperformance liquid chromatography was performed using an Acquity PREMIER BEH C18 UPLC column (2.1 x 150 mm, 1.7 μm particle size) (Waters Corp., Mildford, MA, USA). The mobile phase consisted of 0.1% formic acid in water (phase A), and 0.1% formic acid in acetonitrile (phase B). The solvent gradient program was as follows: 0.5% phase B over the first 2 min, 0.5–30% phase B over 25 min, 30–100% phase B over 13 min, 2 min at 100% B, return to the initial 0.5% phase B over 1 min, and conditioning at 0.5% B for 2 minutes. The flow rate was 0.4 ml/min and the column temperature was set at 40 °C. Ionisation was performed with heated electrospray ionization (H-ESI) in negative mode. A standard curve was generated with authentic standards, using genistein as the internal standard. The TraceFinder software (Thermo Scientific, Waltham, MA, USA) was used for data processing. The data were normalized to the mean of the WT samples.

### Expression analyses by RT-qPCR

To examine *SMR1* expression levels, total RNA was extracted from approximately 100 mg of ground tissue using the NucleoSpin RNA Plant Kit (Macherey-Nagel) and treated with DNase I on the column, according to the manufacturer’s instructions. cDNA was prepared from 1.5 μg of total RNA using the PrimeScript 1st strand cDNA Synthesis Kit (Takara). PCR reactions were performed with 1 μl of a 1:4 dilution of cDNA in ddH_2_O, SYBR Premix Ex Taq II (Takara) and the primers listed in Supplementary Table 1; using a 7500 Fast Real-Time PCR System and the associated 7500/7500 software (Applied Biosystems). Relative expression levels were calculated according to the 2^-ΔΔCt^ method ^55^, using the *At1g13320* (*PDF2*) gene as a reference. All experiments included three biological replicates, each measured as the mean of three technical replicates. For the analysis of WT and *gai-1* shoot apices, the biological replicates included five to six inflorescence tips (5 mm in length) with floral buds beyond stage 12 removed ^56^. For the heat shock assays with WT and *pHS:gai-1* plants, the biological replicates contained approximately 50 seedlings grown on plates for 10 days. These assays included *SCL3* (*At1g50420*) as a positive control for DELLA activity ^57^.

The expression levels of *CHS* and *FLS1* were examined in the same plant material used for the quantification of flavonols. Total RNA extraction, cDNA preparation and PCR were performed as described above. Gene expression was normalized to the geometric mean of the *PDF2* and *TIP41* (*At4g34270*) standards. Five to six plants (biological replicates) were analyzed in triplicate per genotype.

### Chromatin immunoprecipitation (ChIP-qPCR)

1-week-old seedlings of *pGAI:gai-1-3xYPet*, *pRGA:GFP-rga-Δ17* and non-transgenic L*er* WT lines grown under continuous light were harvested. In vitro double cross-linking with ethylene glycol bis(succinimidyl succinate) (EGS) and formaldehyde, chromatin extraction, chromatin sonication, chromatin immunoprecipitation with an anti-GFP antibody (5 μg of ab290 per sample; Abcam), and DNA purification were performed as previously described ^43^. ChIP-qPCR reactions were carried out in the same manner as qPCR analyses using the primers listed in Supplementary Table 1. In addition to the *SMR1* genomic regions, a fragment of the *SCL3* promoter was amplified as a positive control for DELLA binding^58^. Enrichments were determined as the ratio of immunoprecipitated DNA to input DNA.

### Confocal imaging of root tips

Four days-old seedlings grown in vertical plates under our standard conditions were transferred to new plates containing fresh ½ MS medium (mock samples) or ½ MS medium containing 20 μg/ml zeocin (Invitrogen) or 3 μM methyl viologen dichloride hydrate (MV, Sigma-Aldrich). Treatments were performed for 24 h under the same growth conditions, unless the zeocin plates and corresponding mocks were covered with foil. For imaging, roots were mounted on slides with 10 μg/ml propidium iodide solution (PI, Sigma-Aldrich) to simultaneously stain cell borders and cell death regions. Confocal longitudinal stacks through the center of the primary root were acquired using a Zeiss AxioObserver 780 confocal microscope with a Plan-Apochromat 20x/0.8 M27 objective. The excitation wavelength of the laser was 561 nm, and the PI fluorescence was recorded in the range of 600-656 nm. The laser power and digital gain were individually adjusted to maximise signal intensity without saturation. The areas of cell death were quantified with Fiji ImageJ in the region between the quiescent center of the root and the elongation zone (defined at the cortex cell layer). A brightness threshold of 160-255 was set to delimit the areas of cell death.

### Confocal imaging of inflorescence tips

For combined labelling with modified pseudo-Schiff-propidium iodide (mPS-PI) and 5-ethynyl-2’-deoxyuridine (EdU), shoot apices from plants with approximatelly three open flowers were cut and dissected to expose the SAM region. The dissected apices were cultured in sterile plastic boxes containing ½ MS medium for 22 h, transferred to new boxes containing ½ MS medium supplemented with 10 μM EdU (Invitrogen) and cultured for 2 h to achieve a recovery time of 24 h. Shoot apices were fixed, stained and processed for imaging as previously described ^59^. Confocal z-stacks (0.5 μm step size) were acquired using a Leica Stellaris 8 FALCON confocal microscope with a HC PL APO CS2 40x/1.25 GLYC objective. Excitation was performed at 500 nm with a white light laser and the emission filters were set to 510-530 nm for EdU and 600-700 for PI. EdU-labelled cells in the SAM L1-L3 cell layers were counted manually using the Multi-point Tool of Fiji ImageJ.

To determine the presence of cell death in zeocin-treated shoot apices, dissected inflorescence tips from plants with approximately three open flowers were grown for 48 h in sterile plastic boxes containing ½ MS medium (mock samples) or ½ MS medium supplemented with 100 μg/ml zeocin. The medium was also supplemented with 300 μg/ml carbenicillin to minimize bacterial growth. Before imaging, shoot apices were stained with 1 mg/ml PI for 5 min. Confocal z-stacks (0.5 μm) were imaged using a Leica Stellaris 8 confocal microscope with a Fluotar VISIR 25x/0.95 water dipping lense. Excitation was at 535 nm and PI fluorescence was recorded at 575-700 nm. For shoot apices expressing a *pCLV3:GFP(-ER)* reporter, excitation was set to 488 nm and an additional emission filter was set to 500-525 nm to collect GFP fluorescence. The laser power and digital gain were adjusted for each sample to achieve the highest signal intensity without saturation. 3D cell death regions in the SAM domain were selected using a brightness threshold of 220-255 and quantified with the Voxel Volume macro of Fiji.

To detect cell death in the SAM region of plants exposed to drought stress, shoot apices from WT and *della^KO^* plants were dissected and immediately stained with PI. Image acquisition and analysis were performed as described above. To check DNA fragmentation, TUNEL assays were performed using the In Situ Cell Death Detection Kit Fluorescein (Roche), following the manufacturer’s protocol with some modifications. Dissected inflorescence tips were fixed with 4% (v/v) paraformaldehyde and 0.1% Triton X-100 in phosphate-buffered saline (PBS) under vacuum for 60 min. After washing with PBS, the samples were treated with a permeabilization solution [3% (v/v) Nonidet P40 (Sigma-Aldrich) and 10% DMSO in PBS] for 60 min at room temperature. After washing with PBS, the samples were incubated with TUNEL reaction mixture for 1 h at 37 °C in a wet chamber. The shoot apices were washed again with PBS and water before imaging. Confocal z-stacks (1 μm step size) were imaged using a Zeiss AxioObserver 780 confocal microscope equipped with a C-Apochromat 40x/1.20 water dipping lense. Fluorescein was excited with a laser line of 488 nm, and emission was detected in the range of 515-565 nm. The imaging parameters were kept constant for all samples.

## Data availability

The authors declare that all data supporting the findings of this study are included in the manuscript and its supplementary information files. The generated plasmids and mutant lines are available from the corresponding authors upon reasonable request.

## Acknowledgements

We thank Karthikbabu Kannivadi Ramakanth in Jian Xu lab (University of Singapore) for advice on TUNEL labelling. Flavonols quantification was performed by Ana Espinosa at the IBMCP Metabolomics Platform. We also appreciate the assistance of Marisol Gascón at the IBMCP Microscopy Service. We thank Cristina Ferrándiz for critical reading of the manuscript and the members of the Plasticity Lab for stimulating discussions (https://plasticity.webs.upv.es). This research has been carried out thanks to grants PID2019-109925GB-I00 (to D.A.), PID2022-141770NB (to M.A.B.) and PID2022-137859OB-I00 (to A.S-M.) funded by the Spanish MICIN/AEI/10.13039/501100011033/ and by “ERDF, A way of making Europe”. A.S-M was also funded by the EU-MSCA program (H2020-MSCA-IF-2016-746396) and the Generalitat Valenciana Plan GenT program (CISEJI/2022/28). Research in R.S. group was funded by BSRC grant BB/X01102X/1 and UKRI Frontier Research Grant EP/X034550/1. Research in A.G-C. group was funded by the Generalitat Valenciana grant CIAICO/2021/063.

## Author contributions

A.S.M, J.H.G., D.A. and M.A.B. conceptualized the study. A.S.M. and J.H.G. performed most experiments. C.d.O. and A.G.C. designed and performed the drought tolerance assays. N.B.T. carried out the ChIP-qPCR analysis. A.S.M., J.H.G., C.d.O., N.B.T., R.S., D.A. and M.A.B. analyzed and discussed data. A.S.M. and M.A.B. supervised the study. A.S.M., J.H.G. and M.A.B. wrote the manuscript with inputs from all authors.

---

**Supplementary Fig. 1.**
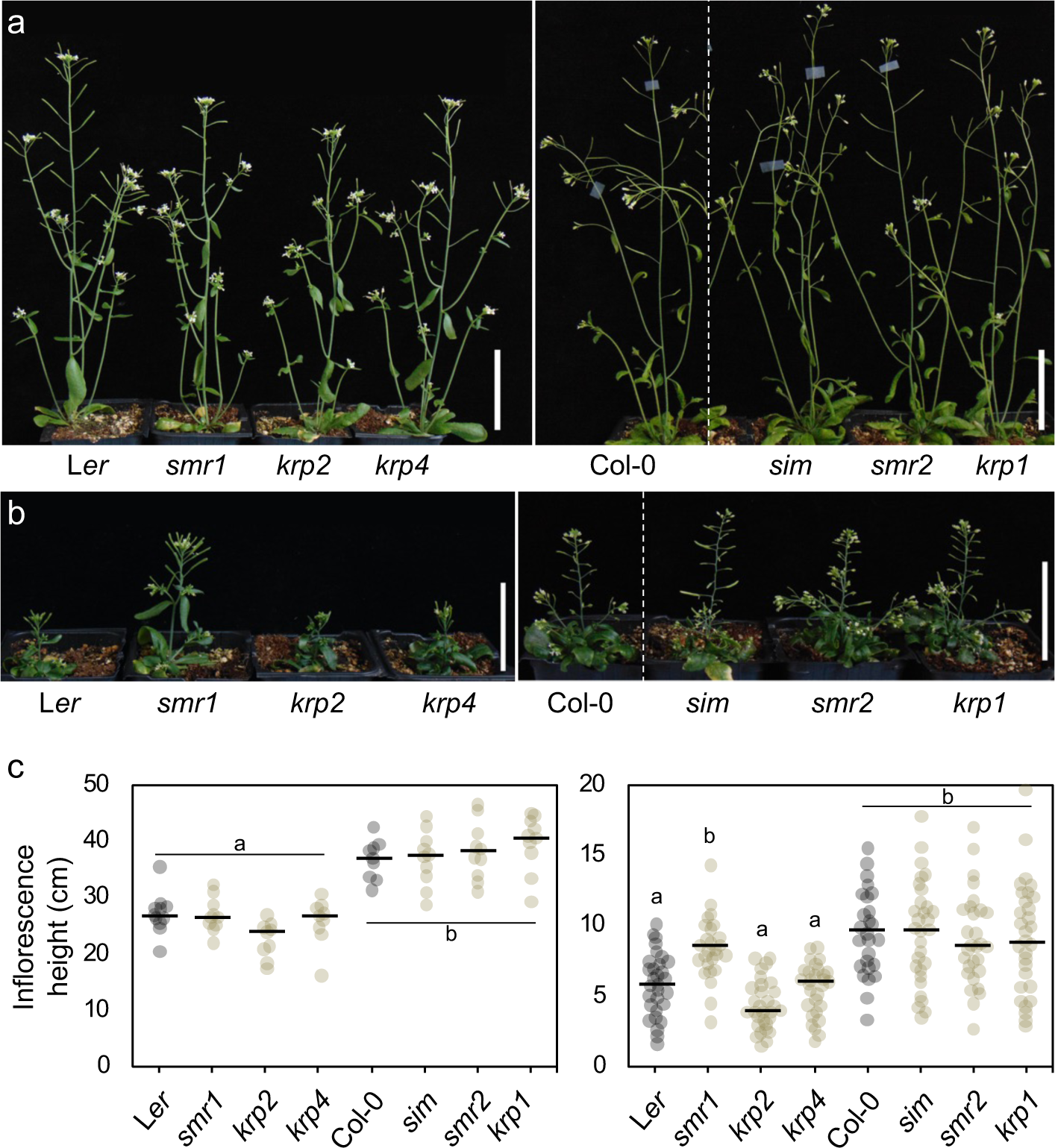
*smr1* shows enhanced resistance to paclobutrazol (PAC) treatment. **a-b** Representative inflorescences of five weeks old plants watered from one week after germination with mock solution (**a**) or 1 μM PAC (**b**). Left panels, loss-of-function mutants in the L*er* background (*smr1-1*, *krp2-3* and *krp4-2*). Right panels, loss-of-function mutants in the Col background (*sim-1*, *smr2-1* and *krp1-1*). **c** Stem length of the main inflorescence of five weeks-old plants watered with mock (graph on the left) or PAC (graph on the right), n ≥ 9 for mock plants and n > 25 for PAC-treated plants. Letters denote statistical groups determined by Tukey’s post hoc test (p < 0.05) after one-way ANOVA. Scale bars, 5 cm.

**Supplementary Fig. 2.**
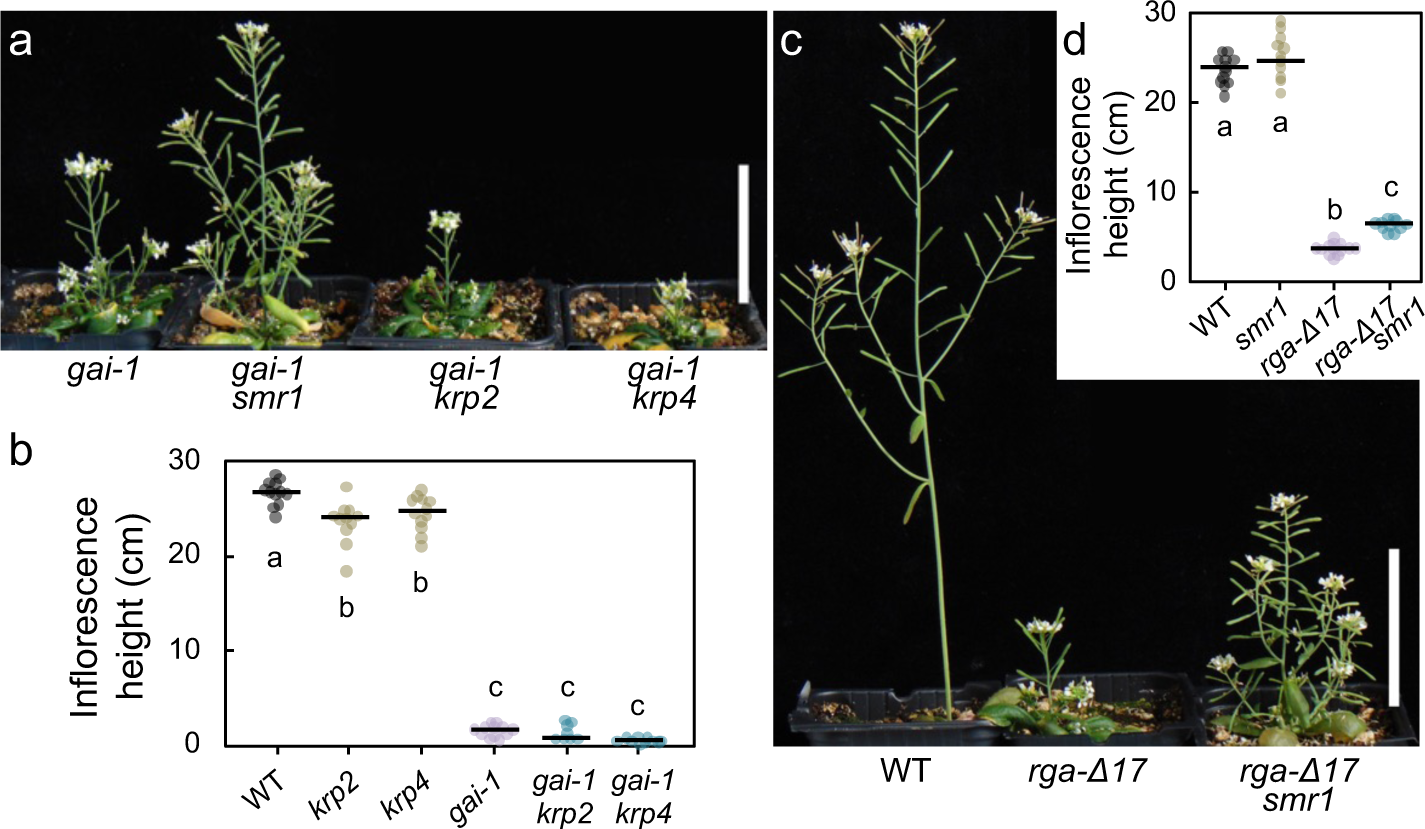
*smr1* promotes the inflorescence growth of *della* gain-of-function mutants. **a** Representative inflorescences of six weeks-old *gai-1*, *gai-1 smr1*, *gai-1 krp2* and *gai-1 krp4* plants. Note that only *smr1* partially suppressed the dwarf architecture of the *gai-1* mutant. **b** Height of the main inflorescence of five weeks-old WT, *krp2*, *krp4, gai-1, gai-1 krp2 and gai-1 krp4* plants. **c** Representative inflorescences of five weeks-old WT, *pRGA:GFP-rga-Δ17 (rga-Δ17)* and *rga-Δ17 smr1* plants. **d** Height of the main inflorescence of five weeks-old WT, *smr1*, *rga-Δ17* and *rga-Δ17 smr1* plants. In (**b**) and (**d**), the dots represent individual values (n = 12) and the horizontal lines represent the mean value for each genotype. Statistical groups were defined using Tukey’s post hoc test [p < 0.01 for (**b**); p < 0.001 for (**d**)] following one-way ANOVA. Scale bars, 5 cm.

**Supplementary Fig. 3.**
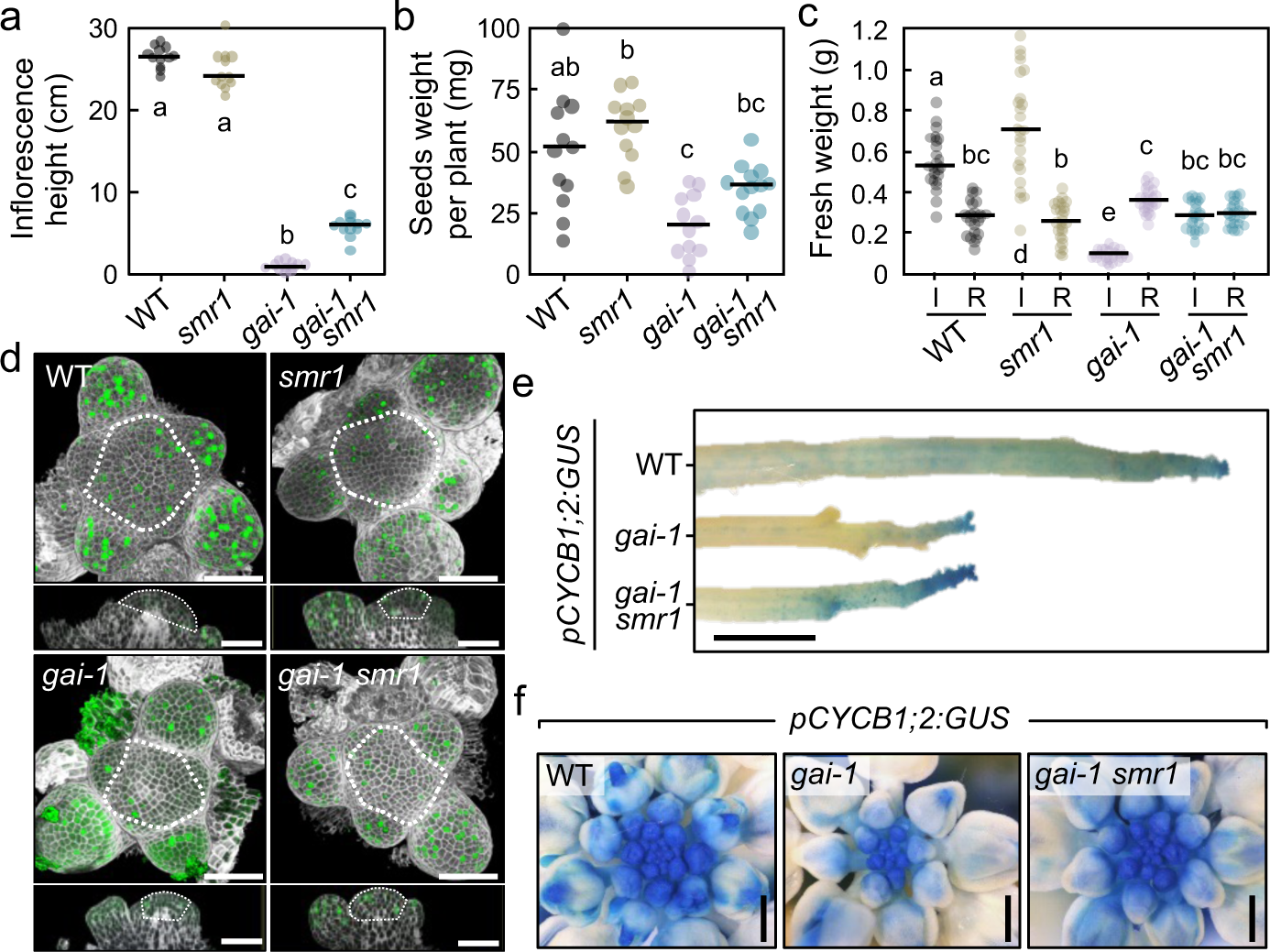
*smr1* restores cell divisions in the shoot apex to promote the reproductive development of *gai-1* mutants. **a-c** Five weeks old inflorescence height (**a**), total weight of seeds (**b**) and aerial biomass (**c**) in WT, *smr1*, *gai-1* and *gai-1 smr1* plants. The dots represent individual plants [n = 12 in (**a**) and (**b**), n = 24 in (**c**)] and the horizontal black lines represent the mean value per genotype. Statistical groups were determined by Tukey’s post hoc test [p < 0.001 for (**a**); p < 0.01 for (**b**); p < 0.05 for (**c**)] following one-way ANOVA. I, total fresh weight of the main inflorescence and secondary inflorescences longer than 0.5 cm. R, total fresh weight of rosette leaves and secondary inflorescences shorter than 0.5 cm. **d** Representative shoot apices stained with mPS-PI to define cell boundaries and labelled with EdU (green signal) to track cell cycle progression. For each genotype, 3D reconstructions from confocal image stacks are shown in the upper panels and longitudinal sections are shown in the lower panels. The dashed lines mark the SAM domain analysed in Fig. 1c. **e-f** Side (**e**) and top views (**f**) of WT, *gai-1* and *gai-1 smr1* inflorescence tips expressing a *pCYCB1;2:GUS* mitotic reporter. The domain of *pCYCB1;2:GUS* activity was reduced in *gai-1* compared with that in WT and *gai-1 smr1*. Scale bars, 50 μm in (**d**), 2 mm in (**e**) and 0.5 mm in (**f**).

**Supplementary Fig. 4.**
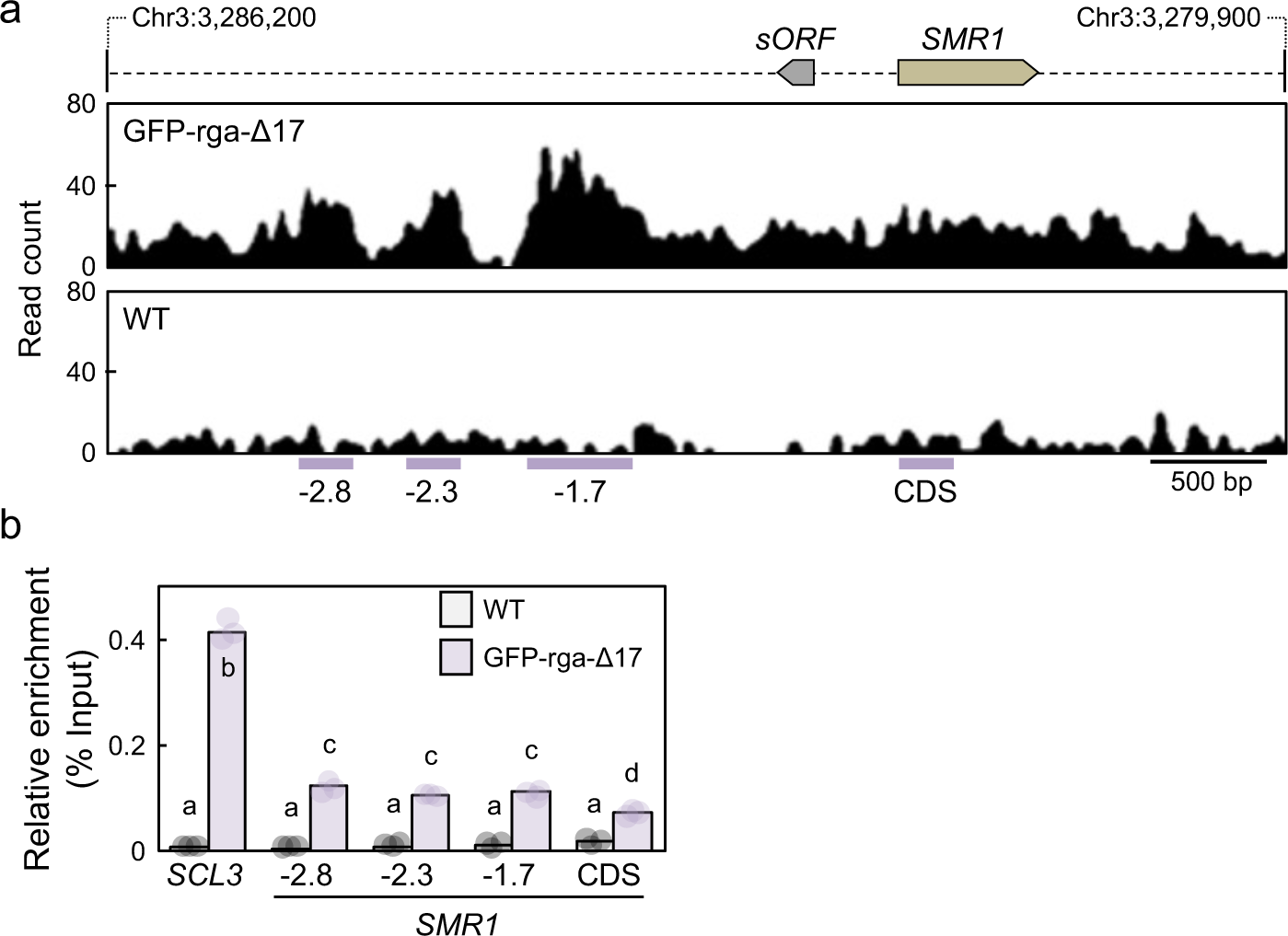
DELLA proteins bind to *SMR1* genomic regions. **a** Representative ChIP-seq peaks of GFP-rga-Δ17 (top graph) and WT control (bottom graph) in the *SMR1* genomic region ^11^. The numbers above the graphs indicate chromosome positions in bp. Purple bars mark the regions that were subsequently analysed by ChIP-qPCR. −2.8, −2.3 and −1.7 indicate the position of these regions in kb relative to the *SMR1* coding sequence (CDS). **b** ChIP-qPCR showing the binding of the GFP-rga-ι117 protein to *SMR1*. Bars represent the mean value of three independent biological replicates (represented by dots). Statistical groups were defined by Tukey’s post hoc test (p < 0.01) following one-way ANOVA.

**Supplementary Fig. 5.**
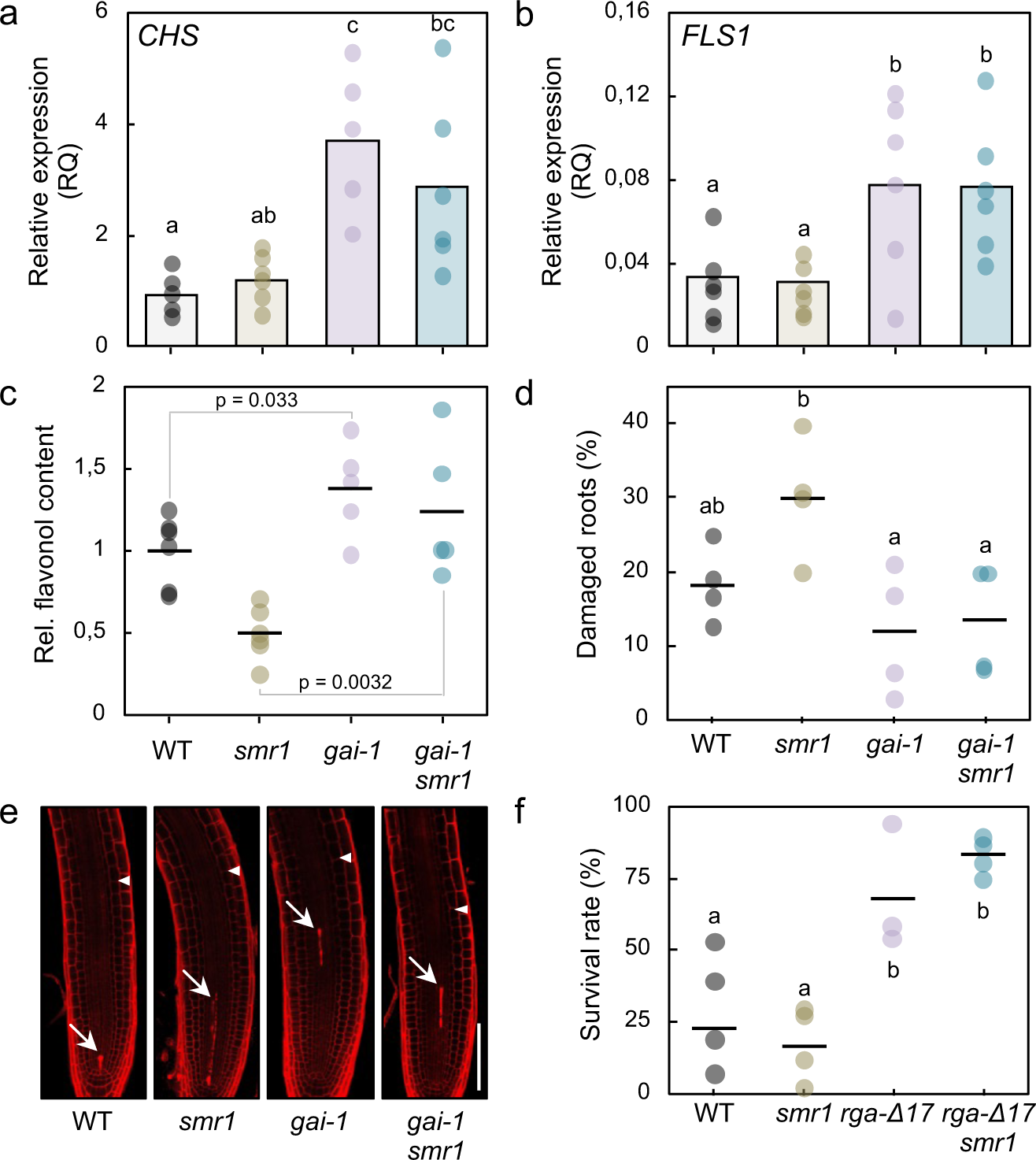
*smr1* has no effect on the antioxidant response of *della* gain-of-function mutants. **a-b** Relative quantity (RQ) of *CHALCONE SYNTHASE* (*CHS*) and *FLAVONOL SYNTHASE1* (*FLS1*) transcripts in WT, *smr1*, *gai-1* and *gai-1 smr1* plants. Bars indicate the mean value of 5-6 plants (dots) per genotype. **c** Relative contents of quercetin and kaempferol in WT, *smr1*, *gai-1* and *gai-1 smr1* plants. Horizontal black lines stand for the mean value of 5-6 plants (dots) per genotype. **d-e** Cell death, detected by PI staining, in root tips treated with 3 μM methyl-viologen for 24 h. In (**d**), the dots indicate the percentage of roots that showed cell death patches in four independent assays (n > 25). Horizontal black lines represent the mean value for each genotype. Representative images are shown in (**e**). Arrows mark dead cells and arrowheads mark the extension of the RAM. **f** Drought resistance of WT, *smr1*, *pRGA:GFP-rga-Δ17 (rga-Δ17)* and *rga-Δ17 smr1* plants. The dots represent independent experiments, with 9-15 plants per genotype, and the horizontal lines represent the mean value for each genotype. The assay was performed as described in Fig. 2d. In (**a**), (**b**), (**d**) and (**f**), letters denote statistical groups that were determined using Tukey’s post hoc test (p < 0.05) following one-way ANOVA. In (**c**), *p* values are for the equality of means (two-tailed *t* test). Scale bars, 50 μm.

**Supplementary Fig. 6.**
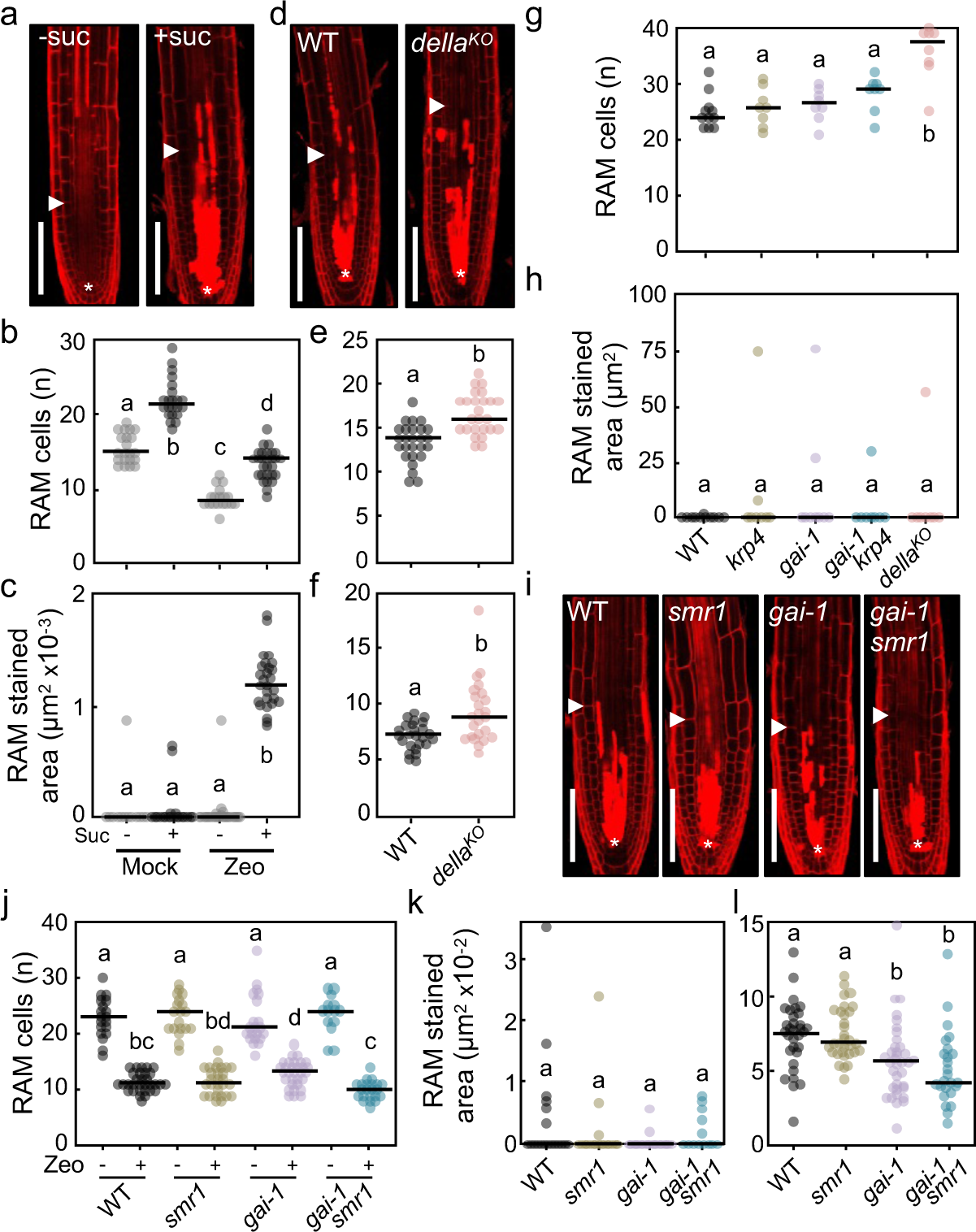
Cell divisions increase the sensitivity of RAM cells to genotoxic stress. **a** Representative confocal images of WT root tips treated with 20 μg/ml zeocin for 24 h in the absence (-suc) or presence (+suc) of 1% sugar. Cell death was detected by PI staining. **b-c** Number of cells in the cortex layer (**b**) and quantification of cell death (**c**) in the central region of the RAM of WT seedlings treated with mock or zeocin (Mock/Zeo) in the absence or presence of sugar (Suc-/Suc+). **d** Representative confocal images of WT and *della^KO^* root tips after incubation in zeocin. **e-f** Number of cortex cells (**e**) and quantification of cell death (**f**) in the central region of the RAM of WT and *della^KO^* zeocin-treated seedlings. **g-h** Number of cells in the cortex layer (**g**) and presence of cell death (**h**) in the central region of the RAM of WT, *krp4*, *gai-1*, *gai-1 krp4* and *della^KO^* seedlings grown under mock conditions. **i** Representative images of WT, *smr1*, *gai-1* and *gai-1 smr1* root tips treated with zeocin. **j** Number of cortex cells in the RAM of WT, *smr1*, *gai-1* and *gai-1 smr1* seedlings treated with mock (Zeo-) or zeocin (Zeo+). **k-l** Quantification of cell death in the RAM of WT, *smr1*, *gai-1* and *gai-1 smr1* seedlings treated with mock (**k**) or zeocin (**l**). In the images of root tips, asterisks mark the position of the quiescent center and arrowheads show the extent of the RAM. In the graphs, the dots represent individual observations [n > 22 in (**b, c**), n ≥ 25 in (**e, f**), n > 7 in (**g, h**), n > 14 in (**j**) Zeo- and (**k**) and n > 25 in (**j**) Zeo+ and (**l**)] and the black horizontal lines represent the mean for each genotype. Letters denote statistical groups determined by Tukey’s post hoc test [p < 0.01 for (**b, c**), (**e, f**) and (**g, h**); p < 0.05 for (**j-l**)] after one-way ANOVA. Scale bars, 50 μm.

**Supplementary Fig. 7.**
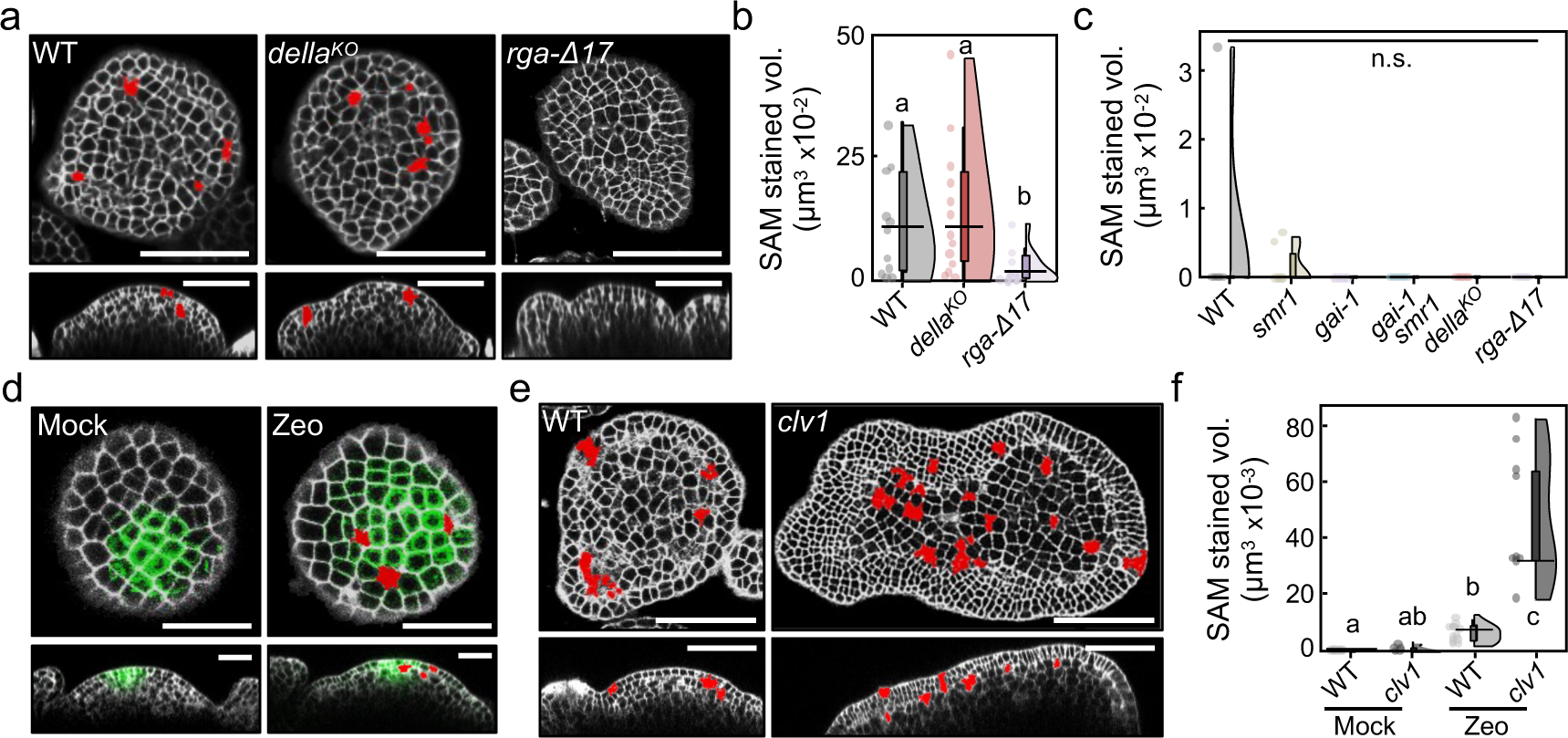
Cell divisions increase the sensitivity of SAM cells to genotoxic stress. **a** Orthogonal views of PI-stained WT, *della^KO^* and *pRGA:GFP-rga-Δ17 (rga-Δ17)* inflorescence apices treated with zeocin. **b** Quantification of cell death in the SAM domain of WT, *della^KO^* and *rga-Δ17* inflorescence tips treated with zeocin. **c** Analysis of cell death presence in the SAM domain of WT, *smr1*, *gai-1*, *gai-1 smr1*, *della^KO^* and *rga-Δ17* inflorescence tips after 48 h incubation in growth medium. **d** Orthogonal views of confocal image stacks of representative WT inflorescence apices expressing a *pCLV3:GFP(-ER)* reporter to mark stem cells (green signal) after 48 h incubation in growth medium (left panels, Mock) or in growth medium containing 100 μg/ml zeocin (right panels, Zeo). PI staining showed that the early descendants of stem cells were more sensitive to the drug. **e** Orthogonal views of representative WT and *clv1* inflorescence apices treated with zeocin. **f** Quantification of cell death in the SAM domain of WT and *clv1* inflorescence tips after incubation in mock or zeocin medium. In (**a**), (**d**) and (**e**), the cell death regions are false colored in red. In (**b**), (**c**) and (**f**), raincloud plots show experimental observations (dots), the median value for each genotype (black horizontal lines) and the distribution of the data (boxplots and split-half violin plots); n > 10 in (**b**), n > 5 in (**c**), n = 5 in (**f**) Mock and n ≥ 9 in (**f**) Zeo. Letters indicate statistical groups determined by Dunn’s post-hoc test following multiple comparisons Kruskal-Wallis (p < 0.05). n.s., not significant. Scale bars, 50 μm in (**a**) and (**e**) and 25 μm in (**d**).

**Supplementary Fig. 8.**
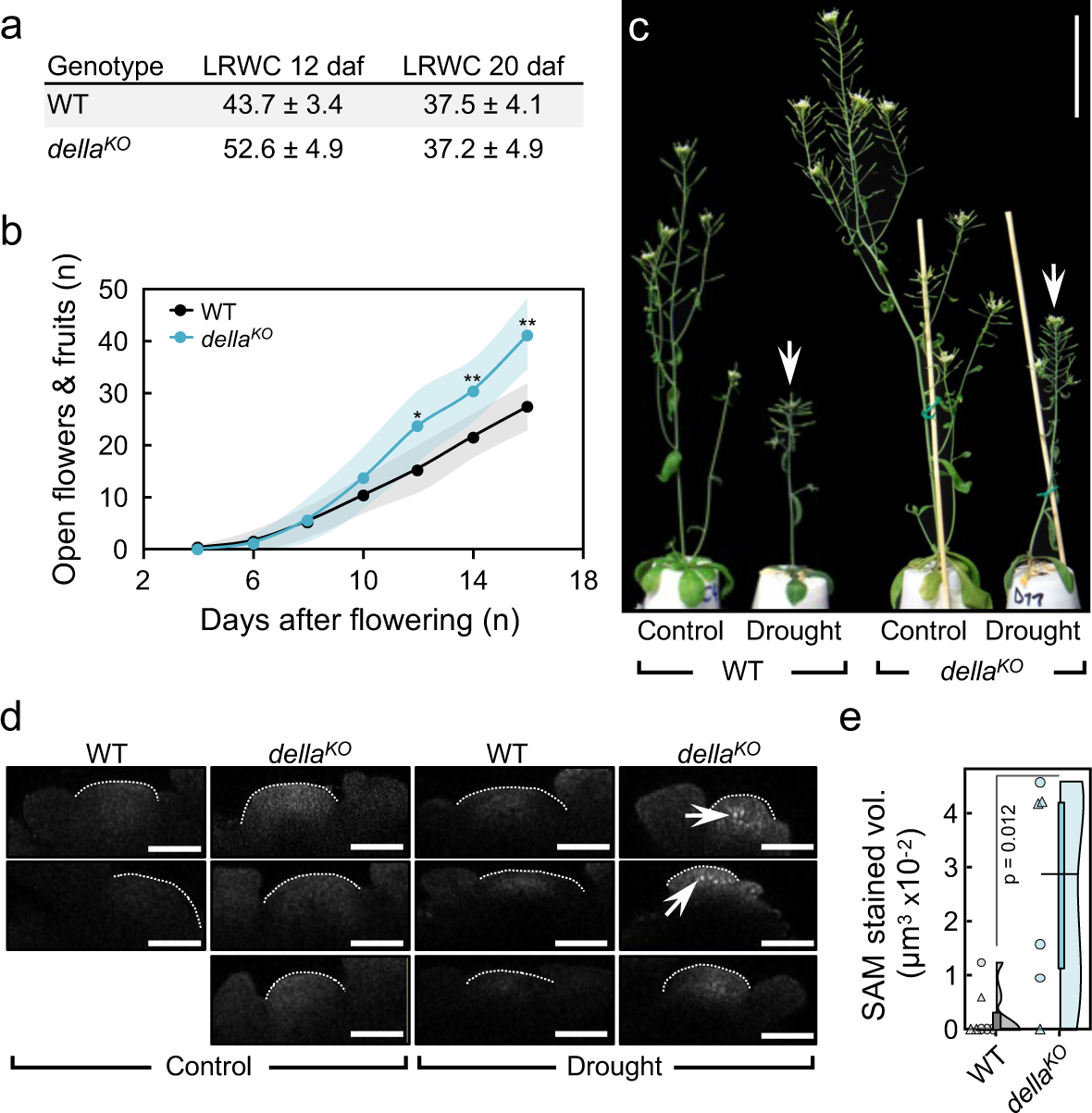
Analysis of SAM activity under drought stress. **a** Relative content of water in leaves (LRWC) of WT and *della^KO^* plants grown under controlled drought conditions for 12 and 20 days after flowering (see related Fig. 4a). The average and standard deviation of four plants per genotype and time point is shown. **b** Cumulative number of mature flowers and fruits in the main inflorescence of WT and *della^KO^* plants grown under well-watered conditions (see related Fig. 4a). The graph shows the mean (dots) and standard deviation (shading) of eight plants per genotype; *p < 0.05, **p < 0.01 (two-tailed *t* test). **c** Representative images of the inflorescence of WT and *della^KO^* plants grown under well-watered conditions (Control) or water restriction (Drought) for 20 days after flowering. Arrows point to the inflorescence apex. Note that that the WT SAM arrested and stopped producing new flowers under drought stress. **d** TUNEL labelling in longitudinal confocal sections through the SAM domain of WT and *della^KO^*plants grown under control or drought conditions. Fluorescence was detected in the SAM of *della^KO^* plants grown under stress (marked with arrows). White dashed lines draw the SAM dome. Panels in the upper row correspond to the shoot apices shown in Fig. 4b. **e** Quantification of cell death in the SAM region of WT and *della^KO^* plants grown under drought stress for three weeks after flowering. Raincloud plots show raw data (n ≥ 6 from two independent experiments represented by circles and triangles), median values (black horizontal lines) and data distribution (boxplots and split-half violin plots); *p* value corresponds to one-tailed Mann-Whitney U test. Scale bars, 5 cm in (**c**) and 50 μm in (**d**).

